# Optoretinography: optical measurements of human cone and rod photoreceptor responses to light

**DOI:** 10.1101/760306

**Authors:** Mehdi Azimipour, Denise Valente, Kari V. Vienola, John S. Werner, Robert J. Zawadzki, Ravi S. Jonnal

## Abstract

Rods contribute crucially to human vision and their dysfunction precedes cones’ in several retinal diseases. Here we describe light-evoked, functional responses of human rods and cones, measured noninvasively using adaptive optics optical coherence tomography.

---

Imaging the stimulus-evoked neural responses of human photoreceptors is an emerging field, with compelling potential applications in basic science, translational research, and clinical management of ophthalmic disease. It is now commonly believed that photoreceptor outer segments (OS) elongate in response to visible stimuli. This biomarker of photoreceptor function has been observed at the cellular level using common path interferometry with adaptive optics (AO) [1, 2]. By providing information about the phase of the light scattered back from single cells, optical coherence tomography (OCT) has been shown capable of detecting elongations much smaller than its axial resolution [3], and this capability has been leveraged to measure light-evoked OS elongation in cones with digital aberration correction [4] and hardware AO [5, 6]. Conventional OCT without AO has been used to observe light-evoked rod elongation in mice [7] and its converse, rod OS shortening during dark adaptation in humans [8]. Functional imaging of single human rod photoreceptors, however, has proven challenging because their small size and rapid functional response place extraordinary demands on the resolution and speed of the imaging system. At present the fastest swept-source systems have wavelengths above 1 μm, which provides insufficient lateral resolution for resolving rods. Alternatives such as full-field OCT may provide sufficient speed but have not yet been used to image rods.

In this work, we employed a combined AO-SLO-OCT to detect light-evoked elongation of rod photoreceptors. The superior lateral resolution afforded by the SLO’s shorter wavelength and sub-Airy disk pinhole was used to confirm the location and type of photoreceptors in the OCT volume. The details of the combined system can be found elsewhere [9]. Briefly, the OCT system was based on a Fourier-domain mode-locked (FDML) swept-source laser with an A-scan rate of 1.64MHz [10]. Three 50:50 fiber couplers in a form of Michelson interferometer was used to split the light between the sample and reference arm. A balanced detector was employed to record the interference pattern and the measured sensitivity of the OCT system was −85 dB. The SLO images were acquired with a superluminescent diode (SLD) (λ = 840 nm; Δλ = 10 nm). Power measured at the cornea was 1.8 mW (OCT) and 150μW (SLO), below the maximum permissible exposure (MPE) specified by ANSI [11]. The AO system was operating in a closed-loop at a rate of 10 Hz and by measuring and correcting aberrations over a 6.75 mm pupil, it provided a theoretical diffraction-limited lateral resolution of 2.75 μm and 3.2μm for the SLO and OCT imaging channels, respectively.

After obtaining informed consent, two normal subjects, free of known retinal disease were imaged. Each subject’s eye was dilated and cyclopleged with drops of 2.5% phenylephrine and 1 % tropicamide. Subjects were dark-adapted for 30 minutes and imaged for 10-15 s, with a 10 ms flash of 555 nm light delivered at 2s with power of either 1 μW, 20 μW, or 80 μW. The flashes were designed to bleach 0.2%, 4%, and 15% of L/M-cone pigment, and 0.05%, 1%, and 4% of rod pigment, respectively. All procedures were in accordance with the tenets of the Declaration of Helsinki and were approved by the University of California, Davis Institutional Review Board. Acquired SLO frames were registered using a strip-based approach, and the resulting trace of eye movements was used to register the simultaneously acquired OCT volumes. Cones and rods were automatically identified in the registered images and axially segmented in the OCT volumes, providing 3D tracking of photoreceptors over time. Time-series of the complex axial signal (M-scans) of each photoreceptor were recorded and the phase difference between the IS/OS and COST and also IS/OS and ROST was measured as a function of time [3, 5].

In Fig. 1, panels (A) and (B) show the OCT en *face* image and SLO frame acquired simultaneously at 6° temporal to the foveal center in a healthy subject. Clusters of rods and some individual rods are visible in (A), distinguishable from cones by their smaller diameters. The rod clusters are revealed as individual rods in (B). The light-evoked elongation of rods and cones for three different bleaching levels are shown in Fig. 2. Panel (A) shows the OCT *en face* image and the locations of selected photoreceptors whose corresponding responses are shown in panel (C) for a 1 μW flash. The responses of cones are not visible above the noise, but rods clearly elongate. Panels (B) and (D) show corresponding results of a 20 μW flash, in which cones and rods are both observed to elongate, but with distinctive dynamics; rods exhibit a slower rate of elongation and slower recovery. Panel (H) shows the amplitude of representative M-scans from a cone and a rod, and plots of their axial profiles, indicating their distinct axial morphology.

**Figure 1:**
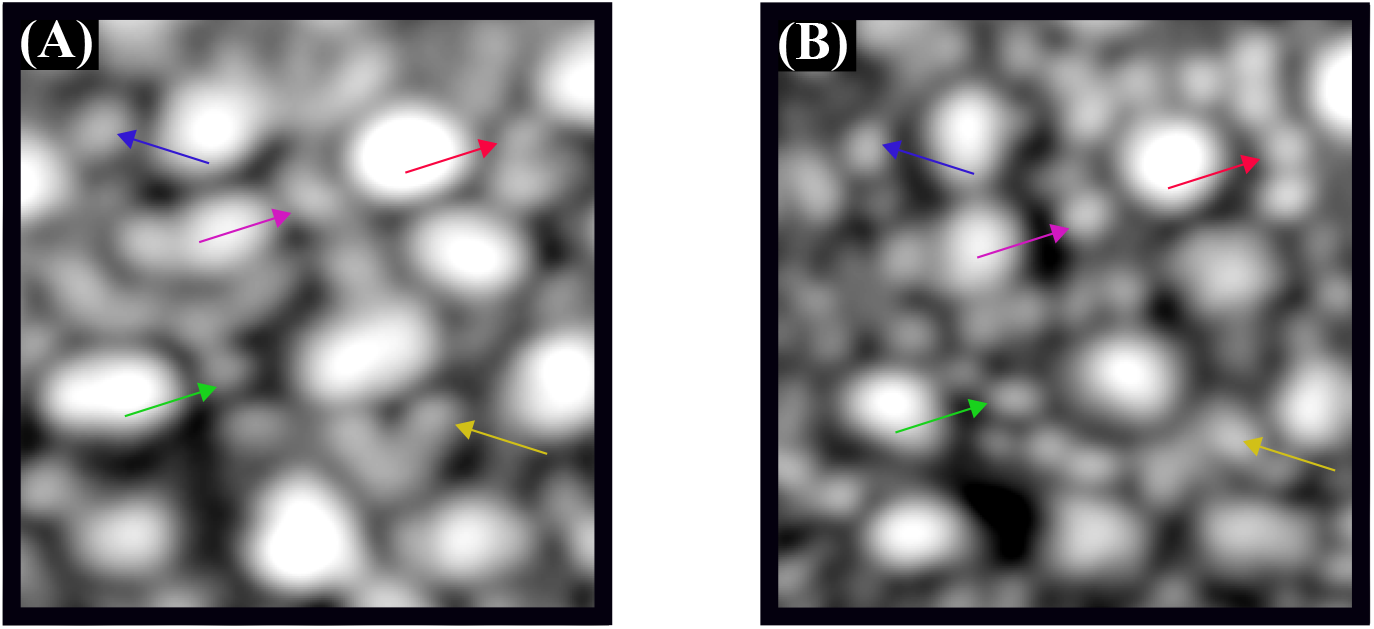
Example of the simultaneously acquired AO-SLO-OCT images at 6° temporal to the foveal center. Rods are not as well resolved in the OCT en *face* projection (A) as they are in the SLO image (B).

**Figure 2:**
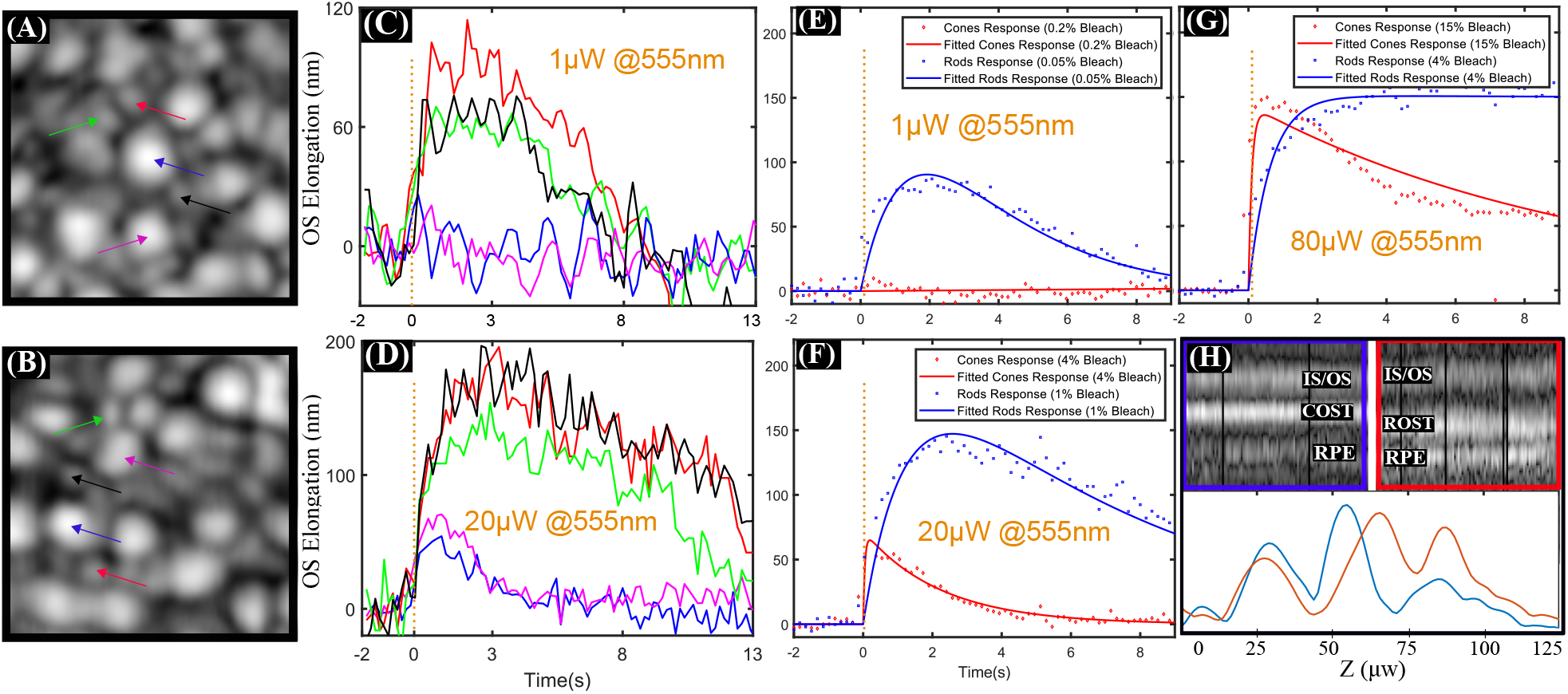
(A) and (B) show the OCT *en face* projections acquired 6° temporal to the fovea. OS elongation of single rods and cones in response to 10 ms flashes of 1 μW and 20 μW, at 555 nm are shown in panels (C) and (D), respectively. No cone response is visible in (C). (E-G) show the averaged response of 10-20 cones and rods to 10 ms flashes with powers of 1 μW, 20 μW and 80 μW, respectively. Rod OS elongation appears to depend on flash energy but saturate at ~ 150 nm. (H) shows a representative M-scan of a cone (top left) and rod (top right) and their axial profiles (bottom), revealing the difference in depth of their OS tips.

This work represents a powerful new tool for noninvasive measurement of both cone and rod photoreceptor function. Such measurements could represent an invaluable new biomarker in the study of retinal disease and development of cures.

## Acknowledgments

The authors gratefully acknowledge the assistance of Susan Garcia and additional funding from the UC Davis Eye Center and a generous donation by Dixie Henderson.

